# The importance of study design for detecting differentially abundant features in high-throughput experiments

**DOI:** 10.1101/007948

**Authors:** Luo Huaien, Li Juntao, Chia Kuan Hui Burton, Paul Robson, Niranjan Nagarajan

## Abstract

The use of high-throughput experiments, such as RNA-seq, to simultaneously identify differentially abundant entities across conditions has become widespread, but the systematic planning of such studies is currently hampered by the lack of general-purpose tools to do so. Here we demonstrate that there is substantial variability in performance across statistical tests, normalization techniques and study conditions, potentially leading to significant wastage of resources and/or missing information in the absence of careful study design. We present a broadly applicable experimental design tool called EDDA, and the first for single-cell RNA-seq, Nanostring and Metagenomic studies, that can be used to i) rationally choose from a panel of statistical tests, ii) measure expected performance for a study and iii) plan experiments to minimize mis-utilization of valuable resources. Using case studies from recent single-cell RNA-seq, Nanostring and Metagenomics studies, we highlight its general utility and, in particular, show a) the ability to correctly model single-cell RNA-seq data and do comparisons with 1/5^th^ the amount of sequencing currently used and b) that the selection of suitable statistical tests strongly impacts the ability to detect biomarkers in Metagenomic studies. Furthermore, we demonstrate that a novel mode-based normalization employed in EDDA uniformly improves in robustness over existing approaches (10-20%) and increases precision to detect differential abundance by up to 140%.

## Introduction

The availability of high-throughput approaches to do counting experiments (e.g. by using DNA sequencing) has enabled scientists in diverse fields (especially in Biology) to simultaneously study a large set of entities (e.g. genes or species) and quantify their relative abundance. These estimates are then compared across replicates and experimental conditions to identify entities whose abundance is significantly altered. One of the most common scenarios for such experiments is in the study of gene expression levels, where sequencing (with protocols such as SAGE^1^, PET^2^ and RNA-Seq^3^) and probe-based approaches^4^ can be used to obtain a digital estimate of transcript abundance in order to identify genes whose expression is altered across biological conditions (e.g. cancer versus normal^5^). Other popular settings where such differential abundance analysis is performed include the study of DNA-binding proteins and histone modifications (e.g. using ChIP-Seq^6, 7^), RNA-binding proteins (e.g. using RIP-Seq^8^ and CLIP-Seq^9^) and the profiling of microbial communities (using 16S rRNA amplicon^10^ and shotgun sequencing^11^).

Due to its generality, a range of software tools have been developed to do differential abundance tests (DATs), often with specific applications in mind, including popular programs such as edgeR^12^, DEseq^6^, Cuffdiff^13, 14^, Metastats^11^, baySeq^15^ and NOISeq^16^. The digital nature of associated data has allowed for several model-based approaches including the use of exact tests (e.g. Fisher’s Exact Test^11^), Poisson^17^ and Negative-Binomial^6, 12^ models as well as Bayesian^15^ and Non-parametric^16^ methods. Recent comparative evaluations of DATs in a few different application settings (e.g. for RNA-Seq^6, 16, 18–21^ and Metagenomics^11^) have further suggested that there is notable variability in their performance, though a consensus on the right DATs to be used remains elusive. In addition, it is not clear, which (if any) of the DATs are broadly applicable across experimental settings despite the generality of the statistical models employed. The interaction between modelling assumptions of a DAT and the application setting, as defined by both **Experimental Choices** (e.g. number of sequencing reads to produce for RNA-seq) as well as intrinsic **Experimental Characteristics** (e.g. number of genes in the organism of interest), could be complex and not predictable *a priori*. Correspondingly, only in very recent work, have experimental design issues been discussed in a limited setting, i.e. using a *t*-test for RNA-seq analysis^22^. Also, as experimental conditions can vary significantly and along several dimensions (Table 1), a systematic assessment of DATs under all conditions is likely to be infeasible. As a result, the choice of DAT as well as decisions related to experimental design (e.g. number of replicates and amount of sequencing) are still guided by rules of thumb and likely to be far from optimal.

**Table 1.** Experimental conditions affecting differential abundance tests (DATs).

|  |  | Abbr. | Description | Notes |
| --- | --- | --- | --- | --- |
| **Experimental Choices** | **Number of Replicates** | NR | Number of technical or non-technical replicates for the two groups compared in the test | For simplicity, in many cases, we assume NR to be the same in both groups |
|  | **Number of Data-points** | ND | Number of data-points generated in the counting experiment | e.g. reads generated in an RNA-seq experiment |
| **Experimental Characteristics** | **Entity Count** | EC | Number of entities in the counting experiment | e.g. number of genes in an RNA-seq experiment |
|  | **Sample Variability** | SV | Variability across replicates (see **Methods**) | e.g. biological variability in RNA-seq datasets |
|  | **Abundance Profile** | AP | Relative abundance of the entities in the first group | Typically follows a power-law distribution |
|  | **Perturbation Profile** | PDA, FC | Perturbations to the abundance profile of the first group to obtain the profile for the second group (see **Methods**) | Used to generate the differentially abundant entities (PDA = Percentage of entities, FC = fold-change distribution) |
| **Test Settings** | **Biases in Data Generation** |  | Deviations from multinomial sampling due to biases inherent in the experimental protocol | These are often corrected for in a pre-processing step e.g. composition-bias in RNA-seq data<sup>35,41</sup> |
|  | **Differential Abundance Test** | DAT | See **Table 2** |  |

In this study, we establish the strong and pervasive impact of experimental design decisions on differential abundance analysis, with implications for study design in diverse disciplines. In particular, we identified data normalization as a source of performance variability and designed a robust alternative (**mode-normalization**) that uniformly improves over existing approaches. We then propose a new paradigm for rational study design based on the ability to model counting experiments in a wide spectrum of applications (Figure 1). The resulting general-purpose tool called **EDDA** (for “Experimental Design in Differential Abundance analysis”), is the first program to enable researchers to design experiments for single-cell RNA-seq, NanoString assays and Metagenomic sequencing and we highlight its use through case studies. EDDA provides researchers access to an array of popular DATs through an intuitive online interface (http://edda.gis.a-star.edu.sg) and answers questions such as “How much sequencing should I be doing?”, “Does the study adequately capture biological variability?” and “Which test should I use to sensitively detect differential abundance in my application setting?”. To provide full access to its functionality, EDDA is also available as a user-friendly R package (on SourceForge: https://sourceforge.net/projects/eddanorm/ and Bioconductor: http://www.bioconductor.org/packages/devel/bioc/html/EDDA.html), and is easily extendable to new DATs and simulation models.

**Figure 1.**
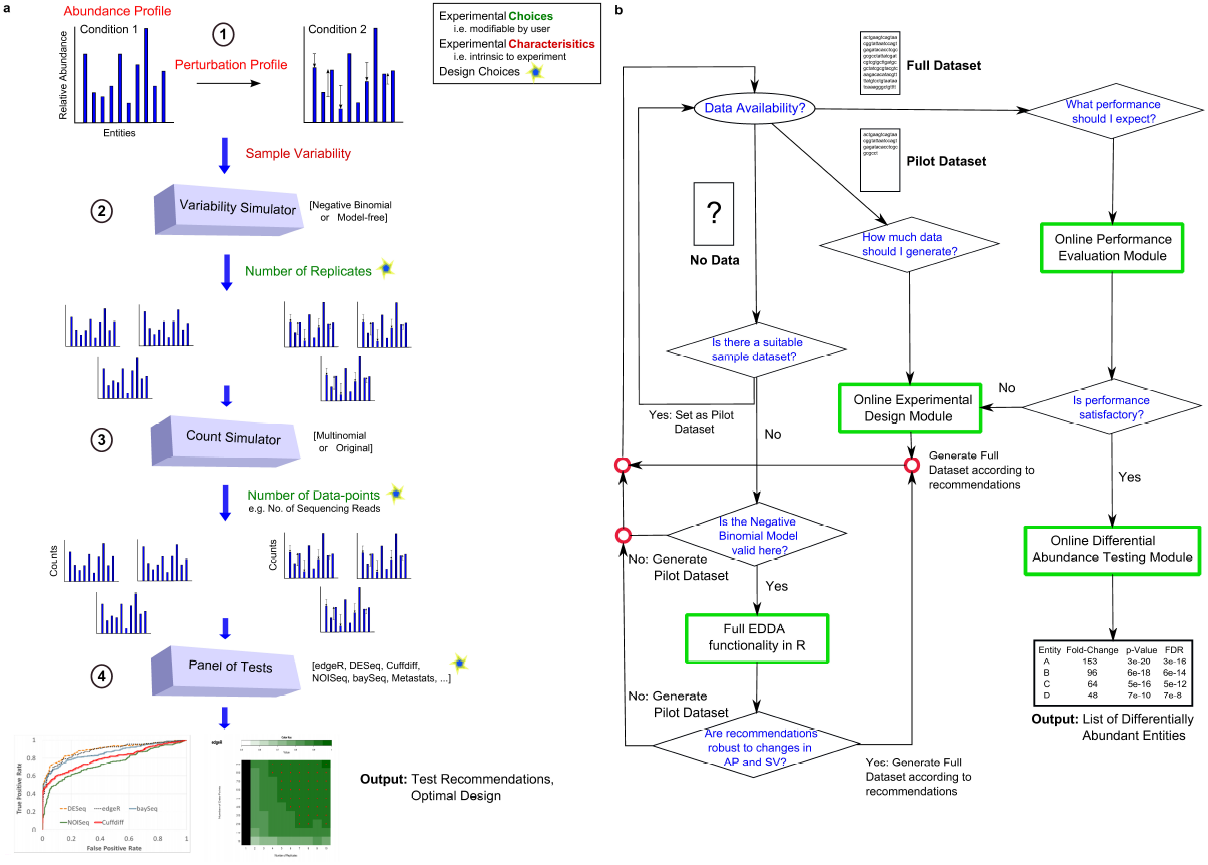
A schematic overview of EDDA and its usage. (a) Overview of the simulate-and-test approach used in EDDA involving 4 key steps (see **Methods** for details). The figure also indicates the various parameters studied here, that impact the ability to predict differentially abundant entities (also see Table 1), in red and green text. (b) A flowchart describing how EDDA can be used to optimize design and performance in a counting experiment. EDDA functionality is depicted in green boxes and rhombi indicate decision nodes.

## Results

In the following, we present and emphasize the somewhat under-appreciated impact of various experimental conditions (grouped into two categories - Experimental Choices and Experimental Characteristics, see Table 1) and various popular DATs (see Table 2) on the ability to detect differential abundance. Our results highlight the importance of careful experimental design and the interplay between experimental conditions and DATs in dictating the success of such experiments. We first establish that the impact of experimental choices on performance can be significant and in the next section explore their interaction with various experimental characteristics. The results presented here are based on synthetic datasets to allow controlled experiments and exploration of a wide-range of parameters, with no emphasis on a particular application. In the following section, we discuss the validity of the modelling assumptions and parameter ranges that we investigated and use this to motivate the design of EDDA. We conclude by showcasing EDDA’s application in various settings. For ease of reference, a schematic overview of the simulation model in EDDA (Figure 1a) and a flowchart of how it can be used (Figure 1b) is provided in Figure 1 with detailed descriptions in the **Methods** section.

**Table 2.** Description of various software packages for conducting differential abundance tests. The various normalization approaches used are: TMM – Trimmed Mean of M values^42^, RPM – Normalization by read count (RNA-seq), RPKM – Normalization by read count and gene length (RNA-seq)^3^, UQN – Upper Quartile Normalization^20^.

| Name | Statistical Testing | Normalization Approach | Target Application Areas |
| --- | --- | --- | --- |
| **edgeR**<sup>12</sup> | Negative Binomial Model, Conditional Maximum Likelihood to estimate parameters, Exact Test | TMM, UQN | SAGE<sup>1</sup>, MPSS<sup>43</sup>, PMAGE<sup>44</sup>, miRAGE<sup>45</sup>, SACO<sup>46</sup> etc. |
| **DESeq**<sup>6</sup> | Negative Binomial Model, local regression to estimate parameters, Exact Test | Normalization by median | RNA-seq<sup>3</sup>, HITS-CLIP<sup>9</sup>, ChIP-seq<sup>47</sup> etc. |
| **baySeq**<sup>15</sup> | Negative Binomial Model, empirical Bayes to estimate parameters | RPM, TMM | DNA-seq, RNA-seq<sup>3</sup>, SAGE<sup>1</sup> etc. |
| **NOISeq**<sup>16</sup> | Non-parametric approach | RPM, RPKM, UQN | ChIP-seq<sup>47</sup>, RNA-seq<sup>3</sup> etc. |
| **Cuffdiff**<sup>14</sup> | *t*-test | RPM, RPKM, UQN | RNA-seq<sup>3</sup> |
| **Metastats**<sup>11</sup> | Non-parametric *t*-test, Fisher's Exact Test for small counts | RPM | Metagenomics |

### Impact of Experimental Choices on Performance

While the availability of high-throughput technologies (such as massively parallel DNA sequencing) to do counting experiments has significantly increased the resolution of such experiments, the cost of the experiment is often still an important factor and the number of replicates and data-points that can be afforded may be less than optimal. Furthermore, in many settings the number of replicates or the number of data-points possible may be constrained due to technological limitations or uniqueness of samples and conditions. In such settings, it would be ideal to understand and exploit the trade-off between the number of replicates and data-points needed (e.g. by doing deeper RNA-seq for a few biological replicates) for such analysis. In the following, we investigate these dimensions individually and in conjunction.

*Number of Replicates* (NR): The performance of DATs (measured here by the area-under-curve of the receiver operating characteristic or ROC curve; AUC) as a function of the number of replicates in the study is highly non-linear, with significant improvements obtainable until a saturation point (indicated by a larger marker size; Figure 2a). The saturation point is likely to be largely determined by the intrinsic variability of replicates in an experiment. However, clear differences are also seen across DATs as seen in Figure 2a, where edgeR and DESeq achieved AUC greater than 0.95 with five replicates, while baySeq required eight replicates and has substantially lower AUC with one replicate. While all DATs seem to converge toward optimal performance (AUC = 1) in this setting, the rate of convergence varies markedly (e.g. note the curve for Cuffdiff). Note that the relative performance of DATs is influenced by the experimental setting, especially in conditions where the number of replicates available is small (as is typically the case), and thus the choice of DAT is strongly dependent on the desired precision/recall trade-off (**Supplementary Figure 1**).

**Figure 2.**
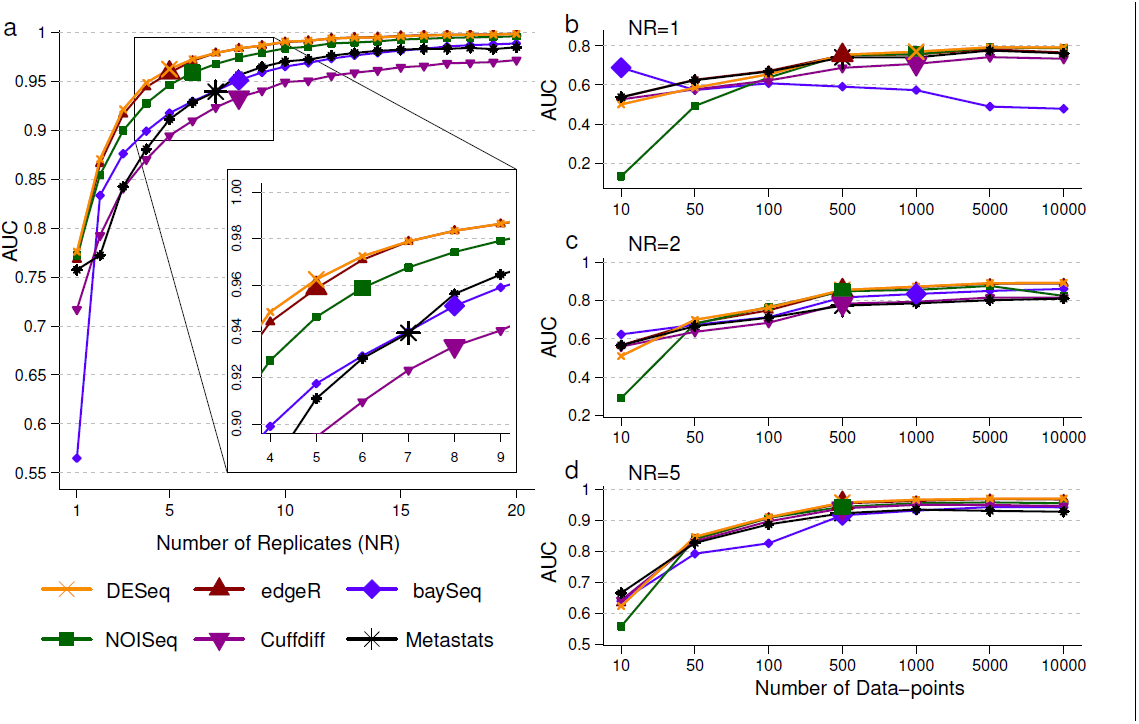
Performance of DATs as a function of Experimental Choices. (a) AUC as a function of the number of replicates. AUC as a function of the number of data-points with (b) 1 replicate (c) 2 replicates and (d) 5 replicates. Note that the number of data-points reported is the average per entity. Points with large markers indicate the saturation point where the AUC is within 5% of the maximum observed AUC. Results reported are based on an average over 10 simulations with the following parameters, EC = 1000, PDA = 26%, FC = Uniform[3, 5], ND = 1000 per entity, AP = BP, SM = NB and SV = 0.85 (see Table 1 and **Methods** for key to abbreviations used here).

*Number of Data-points* (ND): The number of data-points generated in an application setting is often set to be the maximum possible given the resources available. However, this may lead to misallocation of resources as suggested by Figure 2b, 2c and 2d. In this setting, increasing the number of data-points continues to improve AUC over a wide range of values (for most DATs). At very high values AUC saturates, but not necessarily at 1 (Figure 2b, 1c). Oddly, for one of the DATs (baySeq), performance decreases with increase in ND – a feature that is not *a priori* evident from its specification (Figure 2b). However, with more replicates, much fewer data-points are needed to obtain high AUC values, suggesting that this is a better trade-off in this setting (Figure 2c, 2d; not necessarily the case in other settings e.g. when the number of replicates is already high or intrinsic variability across replicates is low). Note that if the number of data-points is limited by the application, then there is considerable variability in performance across DATs (**Supplementary Figure 2**) and some DATs may have consistently lower AUCs (Cuffdiff and Metastats, in this setting) with high ND as well. Thus, to meet experimental objectives, especially when high precision is desired, informed decisions on statistical test and number of replicates to employ are needed (as facilitated by experimental design tools such as EDDA).

### Interaction of Experimental Characteristics and Choices

It is important to note that experimental choices alone do not dictate the ability to detect signals of differential abundance and as we show here, the intrinsic characteristics of the experiment are also an important variable that need to be taken into account. This excludes the possibility of pre-computing recommendations for experimental choices and DATs to use in various applications and emphasizes the need for a general-purpose experimental design tool such as EDDA.

*Entity Count* (EC): Intuitively, the impact of the number of entities being profiled is expected to be minimal and the *a priori* assumption is that scaling the number of data-points as a function of the number of entities should lead to comparable performance. Our results suggest that this is not quite true. As seen in Figure 3a, when the number of entities is low, there is not only greater variability in performance, but average AUC is also lower. Some statistical tests seem to be less appropriate when the number of entities is low (e.g. Cuffdiff), while others exhibited greater robustness (edgeR and DESeq, followed by baySeq).

**Figure 3.**
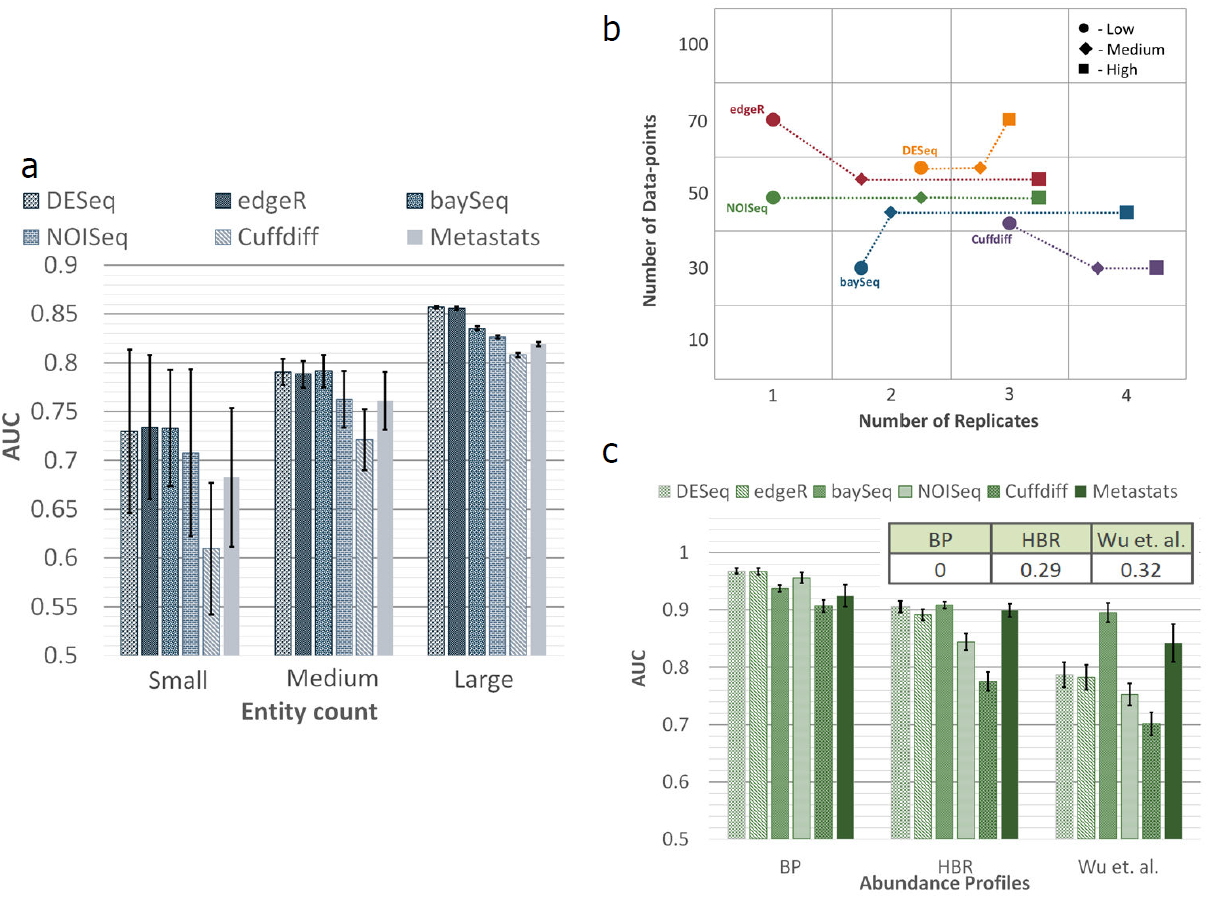
Performance of DATs as a function of Experimental Characteristics. (a) AUC as a function of entity count (small = 100, medium = 1,000, large = 30,000). Simulations were done with the same parameters as in Figure 2a, except with ND = 100 per entity, AP = HBR, SM = Full and FC = Log-normal[2, 1] (b) AUC as a function of SV (and NR, ND per entity), where low, medium and high were set to SV = 0.05, 0.5 and 0.85 respectively. Note that the plotted point shows the smallest NR and ND values (small NR was given priority in the case of ties) that achieved an AUC target of 0.95. Synthetic data was generated with EC = 1000, PDA = 10%, FC = Uniform[8, 8], AP = BP and SM = NB. (c) AUC as a function of abundance profile. Simulations were done with the same parameters as in Figure 2a and with NR = 5 and ND = 500 per entity. The inset shows the fraction of rare entities (mean count < 10) in each abundance profile. Results reported are based on an average over 10 simulations and error bars in subfigures a and c represent one standard deviation.

*Sample Variability* (SV): The intrinsic variability seen across replicates in an application setting dictates the trade-off between the number of data-points and replicates needed in complex ways as shown in Figure 3b. While the specific patterns will depend on the application, even for a setting with large effect sizes as shown here, the specific tradeoffs chosen by the various DATs vary (more sequencing for DESeq vs more replicates for Cuffdiff) and the cost-effectiveness of a method (= NR x ND needed) can switch with sample variability (e.g. Cuffdiff goes from being the least to the most cost-effective when sample variability increases; Figure 3b).

*Abundance profile* (AP): The relative abundance of entities is often seen to follow a power law distribution (**Supplementary Figure 3**), but the precise shape can vary and together with the number of data-points generated, impact overall performance for an application. In particular, testing differential abundance for rare entities (with low relative abundance) can be difficult and could explain the variability in performance seen in Figure 3c. While all methods have lower AUC for a rare-entity-enriched profile from Wu et al^23^ (**Supplementary Figure 3**), some methods seem to be more robust (e.g. baySeq) or tuned to detect rare entities Metastats), while others experience a larger relative drop in performance (e.g. Cuffdiff or NOISeq), suggesting that DAT choices need to take abundance profiles into account.

*Perturbation profile*: The affect of specific profiles of differential abundance on prediction performance is likely to be the least predictable from first-principles and this was also seen in our experiments (Figure 4). Altering the fraction of differentially abundant entities alone could reorder the performance of various statistical tests, as seen in Figures 4a and 4b, where baySeq went from being the worst performer to the best performer. Furthermore, switching the distribution of fold-changes was also seen to affect results as seen in Figures 4b and 4c, with NOISeq now becoming the best performing DAT. Other parameters such as the abundance profile also combine with the perturbation profile to influence relative performance as seen in Figures 4b and 4d, where DEseq went from being one of the best to being the worst performer. Overall, no single DAT was found to outperform others (**Supplementary Figure 4**), highlighting that specific experimental characteristics and choices need to be taken into account while choosing an appropriate DAT.

**Figure 4.**
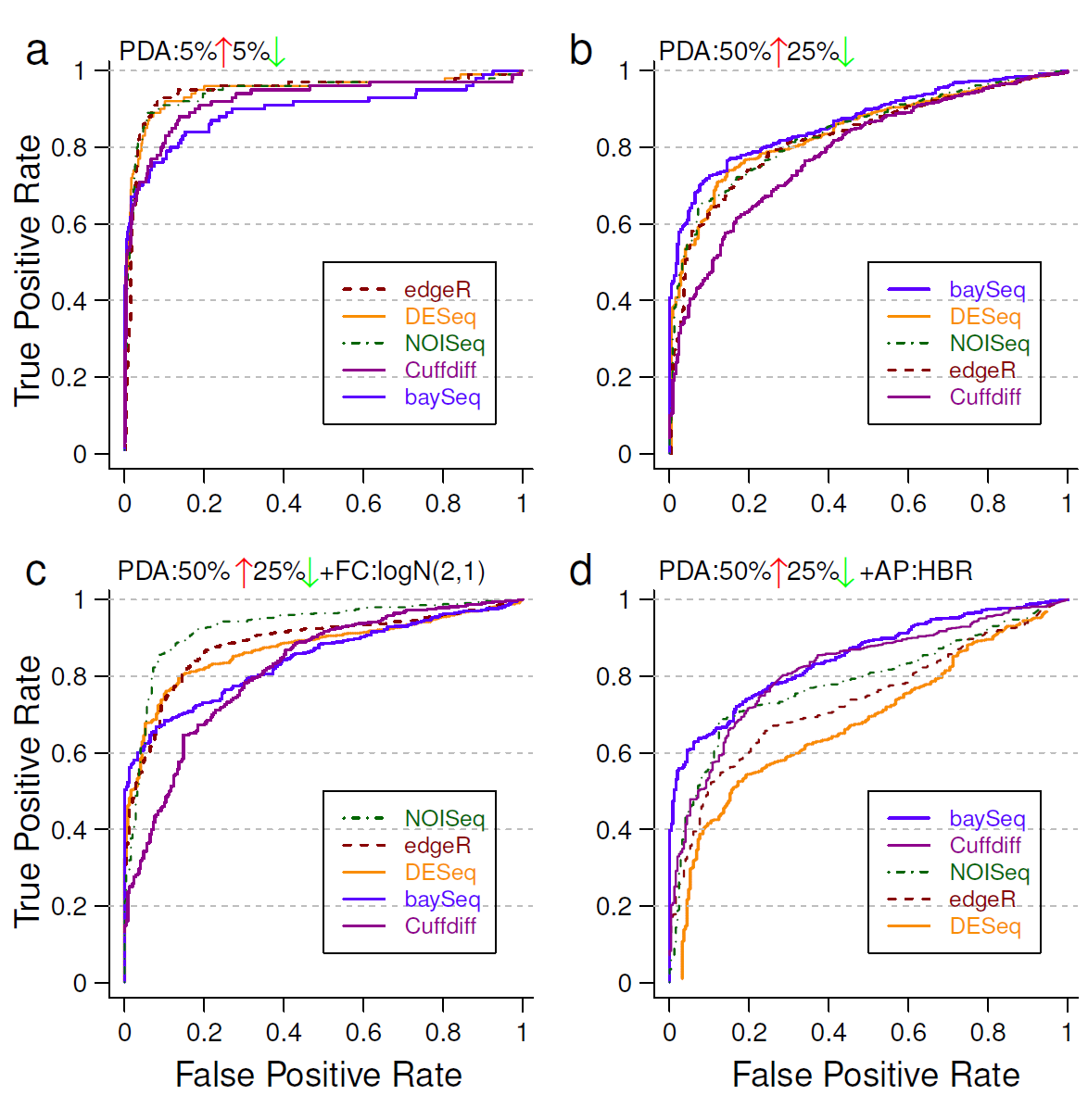
ROC plots under various Perturbation Profiles. Statistical tests in the legend are listed from best to worst (in terms of AUC values) for each setting. Note the striking re-ordering of test performance across subfigures with slight changes in experimental conditions. Simulations in EDDA were done with (a) PDA = 10% (b) PDA = [50% UP, 25% DOWN] (c) PDA = [50% UP, 25% DOWN], FC = Log-normal[2, 1] (d) PDA = [50% UP, 25% DOWN], AP = HBR. Unless stated otherwise, common parameters include NR = 3, EC = 1000, FC = Uniform[3, 7], ND = 500 per entity, AP = BP, SM = Full and SV = 0.5.

### Modelling Assumptions and Normalization

In the absence of variability across replicates and experimental biases, counting experiments of the sort studied here can naturally be modelled as samples from a Multinomial distribution. To simulate technical and intrinsic variability, a common approach has been to model the relative abundance of each entity across replicates using the Negative Binomial distribution^15, 24^. In fact, in many studies this is the model from which counts are simulated for each entity^15, 25^, independent of those for other entities (we refer to this as the **Negative Binomial** model), and bypassing the joint simulation of counts from a Multinomial distribution (referred to here as the **Full** model). In practice, both the Full and the Negative Binomial model elicit similar performance for most experimental settings and most DATs. However, for a few DATs (baySeq, NOISeq and Cuffdiff) we observed deterioration in performance on the Full model when compared to a similar experiment using the Negative Binomial model (**Supplementary Figure 5**) suggesting that the Full model is a better measure of performance of a DAT.

By analysing several published and in-house datasets we established that, in general, for bulk transcriptome sequencing (confirming earlier reports^15, 24^), Nanostring assays and Shotgun Metagenomic sequencing (not shown in prior work), variability in replicates can be adequately modelled using the negative binomial distribution (**Supplementary Figure 6**). An exception to this rule was, however, seen in single-cell RNA-seq experiments in accordance with observations of unusually high cell-to-cell variability in recent reports^26, 27^ (**Supplementary Figure 6c, d**). For cases where an appropriate model for variability across replicates is not available (as in the single-cell case), we developed a **Model-Free** approach, that uses sub-sampling (and appropriate scaling where needed) of existing datasets to provide simulated datasets that match sample-to-sample variability in real datasets better (with the drawback that it relies on the availability of a dataset with many replicates; see **Methods**).

To evaluate the data generation models used in this study (either model-based or model-free), as well as establish their suitability for the design of EDDA, we first investigated distributional properties of real and simulated datasets (**Supplementary Figure 7**). The results here indicate that while overall both simulation approaches (where applicable) provide good approximations and capture the general trend, the Model-Free approach more closely mimics true sample variability (**Supplementary Figure 7**). We next tested the suitability of an approach where simulated datasets are generated to mimic an existing pilot dataset and employed to measure trends in performance. Our results confirmed that simulated data generated by our simulation models enable reliable measurement of true performance for DATs (relative-error in AUC < 6%) and monitoring of trends as a function of experimental choices and characteristics (**Supplementary Figure 8**). In addition, experimental recommendations from EDDA simulations were also found to match DAT recommendations based on benchmarking on real datasets^28^ (**Supplementary Figure 9**), suggesting that EDDA can help avoid this step and still reliably guide experimental design.

In some experimental settings, variability in replicates can be extremely low and directly simulating from the Multinomial distribution (a special case of the Full model that we refer to as the **Multinomial** model; see **Methods**) is sufficient. In principal, with enough data-points, statistical testing under the Multinomial model should be straightforward and we expect various DATs to perform well. The few exceptions that we noted, suggest that aspects other than statistical testing, such as data normalization, may play a role in their reduced performance (**Supplementary Figure 5**).

An investigation of different normalization approaches (Table 2) under the various experimental conditions explored in this study suggests that their robustness can vary significantly as a function of the experimental setting. In particular, we observed a few settings under which many of the existing approaches performed sub-optimally (Figure 5a) and to address this we designed a new method (**mode-normalization**) that analyzes the distribution of un-normalized fold-changes of entities using mode statistics to select a suitable normalization factor (see **Methods** and **Supplementary Figure 10**). We compared mode-normalization to the default normalization and a popular alternate (Upper-quartile Normalization), for each DAT and across all the conditions tested here, to find that the use of mode-normalization uniformly improved performance (on average, AUC by 9% and precision by 14% at 5% FDR). Also, in cases where the performance of a few DATs dipped under the Multinomial model, mode-normalization was able to rescue the AUC values (**Supplementary Figure 5d**). In addition, we identified several examples where mode-normalization significantly improved AUC values for all the DATs tested (improving precision to detect differential abundance by up to 140% at 5% FDR), highlighting that proper data normalization is a key step in attaining experimental goals (Figure 5a). As depicted in Figure 5a, there are often cases where the default normalization of a DAT or a popular alternate (Upper-quartile Normalization) lead to reduced performance while calling differentially abundant entities, while mode normalization consistently achieves optimal performance across DATs. Note that, no normalization can be expected to work under all conditions and simulated datasets generated by EDDA can also be valuable to compare and choose among alternative normalization techniques.

**Figure 5.**
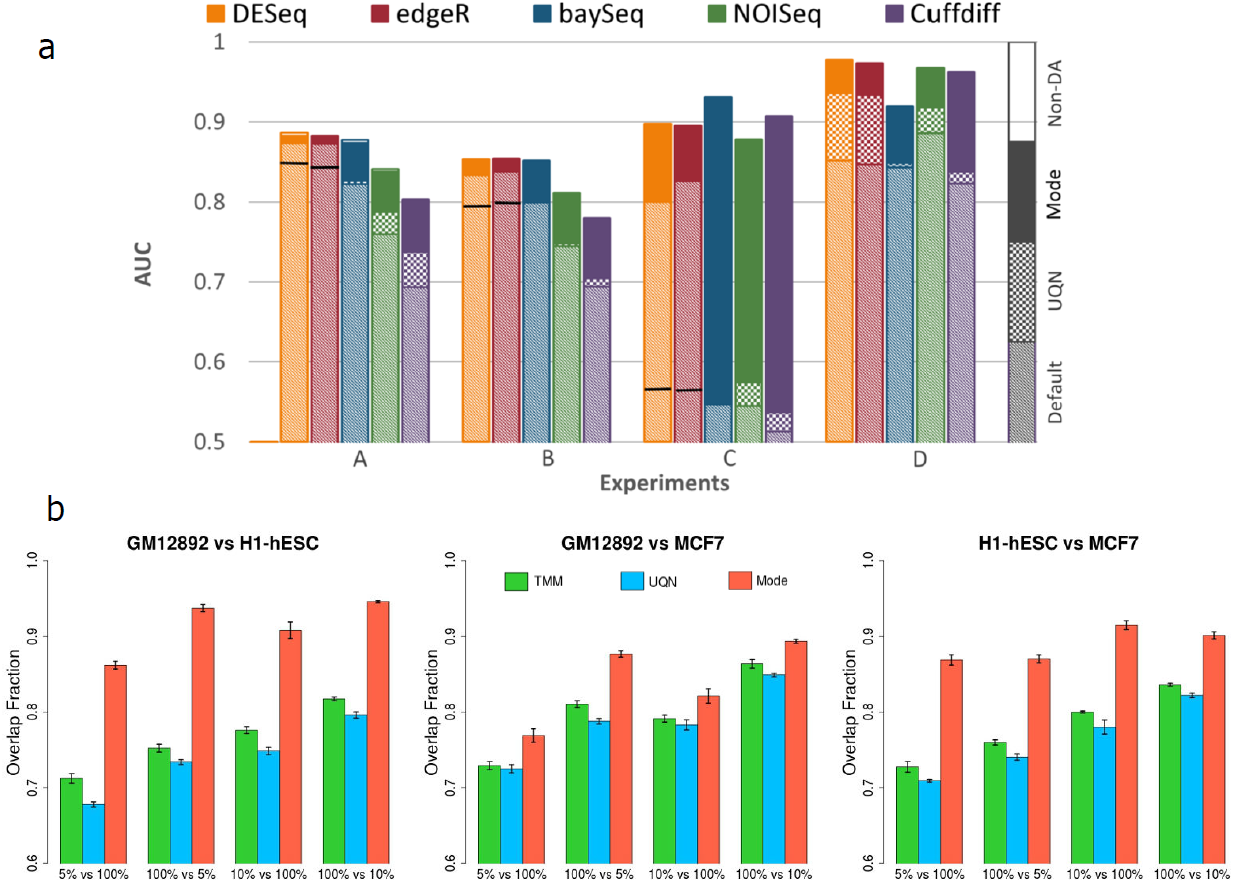
Comparison of different data normalization approaches. (a) On simulated benchmark datasets using EDDA. Note that we used normalization by the sum of counts for all non-differentially-abundant entities (non-DA) as a measure of ideal performance here. In general, Upper Quartile Normalization (UQN) improved over the Default normalization (improvement shown by the checked box) but for cases where it did not we mark its performance with a solid line. Mode normalization always improved over the performance from UQN and the Default normalization for the DAT (improvement shown by the solid box). The parameters for the various experiments include for A: PDA = [26% UP, 10% DOWN], AP = Wu et al B: PDA = [35% UP, 15% DOWN] C: PDA = [40% DOWN],SM = Mutinomial and D: the same as in Figure 4b. Unless stated otherwise, common parameters include NR = 3, EC = 1000, FC = Log-normal[1.5, 1], ND = 500 per entity, AP = HBR, SM = Full and SV = 0.5. (b) Comparison on real datasets, highlighting the robustness of mode normalization. Shown here is the overlap fraction (used to measure robustness) of the top 500 differentially abundant genes (sorted by *p*-value using edgeR) in comparisons involving both the original full-size libraries versus those where one of the libraries is down-sampled (5% or 10% of original size). Barplots show the average of 5 runs and error bars represent one standard deviation.

To further evaluate normalization methods on real datasets, we studied the consistency of differential abundance predictions (against predictions on the full dataset) upon down-sampling of data-points, using 3 deeply-sequenced RNA-seq datasets^28^ (Figure 5b). These results highlight the robustness of mode-normalization versus other popular approaches (UQN and TMM; Table 2). Mode-normalization was found to improve robustness by 10-20% across datasets and was the least affected by imbalances in sequencing depth across conditions (Figure 5b).

### Applications of EDDA and mode-normalization

The observed variability in the performance of DATs across experimental characteristics and choices, and the demonstration that data from many kinds of high-throughput experiments can be adequately modelled *in silico*, motivated the use of a *simulate-and-test* paradigm in EDDA to guide experimental design (see Figure 1a and **Methods**). EDDA allows users fine-scale control of all the variables discussed here (summarized in Table 1), but also provides the option to directly learn experimental parameters (and models for the model-free approach) from pilot or publicly-available datasets. Some of the commonly expected modes of usage for EDDA are discussed in the methods section **EDDA Modules** and illustrated in Figure 1b. Furthermore, to showcase the use of EDDA and mode-normalization, we present results from EDDA analysis of several recently generated datasets in three different experimental settings, each highlighting a different aspect of the utility of the package in a practical scenario.

For the first case study, we analyzed data from a recent single-cell RNA-seq study of circulating tumor cells (CTCs) from melanoma patients^29^. The authors generated on average 1000 data-points per entity (> 20 million reads) and used a one-way ANOVA test (equivalent to a *t*-test) to identify differentially abundant genes between CTCs and primary melanocytes. We reanalyzed the data using EDDA to simulate synthetic datasets that mimicked real data (with the **Model-Free** approach and a 96-cell dataset generated as a resource for this study) and used them to test a panel of DATs (see **Methods**). The availability of new micro-fluidics based systems to automate single-cell omics has highlighted the cost of sequencing as the major bottleneck in studying a large number of cells. Strikingly, EDDA analysis revealed that this study could have been conducted with 1/5^th^ of the sequencing that was done (by reducing sequencing depth to 1/10^th^ and doubling the number of replicates) without affecting performance in terms of identifying differentially abundant genes (Figure 6a). This was, however, only possible if the appropriate DAT was used (edgeR and BaySeq, in this case), with the choice of DAT playing a more significant role than the amount of sequencing done. Using BaySeq with on average 100 reads per gene (i.e. 2 million reads per cell as opposed to the 20 million reads used in this study) and increasing the number of replicates from 5 to 50 (and thus maintaining sequencing cost) would be expected to boost AUC from 0.86 (and 0.75 using the *t*-test) to 0.96 and sensitivity from 57% to 72% at 5% FDR, in this study (Figure 6a). Note that while practical considerations could limit the number of CTCs that can be captured and studied, this should not be an issue for other cell types in this study.

**Figure 6.**
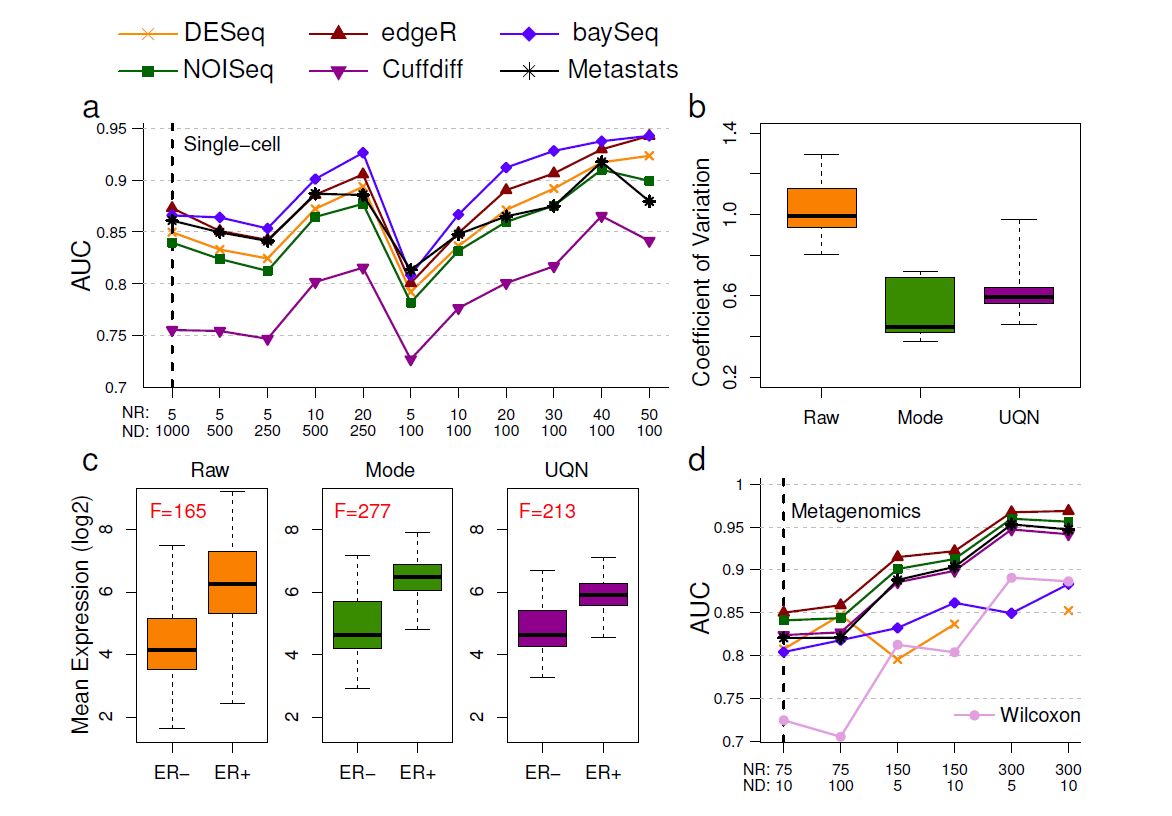
Case studies of EDDA usage. (a) Predicted performance of DATs as a function of replicates (NR) and sequencing depth (ND) for a single cell RNA-seq experiment (Ramskold et al^29^). Note that the vertical black dashed line indicates design choices in the original study. (b) Coefficient of variation for a panel of housekeeping genes from a NanoString experiment under different normalization approaches. (c) Separability of ER positive and negative samples under different normalization approaches. (d) Predicted performance of DATs as a function of replicates (NR) and sequencing depth (ND) for a Metagenomics experiment (Qin et al^31^). Note that the vertical black dashed line indicates design choices in the original study.

In the second case study, we analyzed data from an in-house project (manuscript in preparation) for the development of prognostic and predictive gene signatures in breast cancer on the NanoString nCounter platform (NanoString, WA, USA). The NanoString platform allows for the digital measurement of gene expression, similar to RNA-seq, but is typically used to profile a small, selected set of genes in a large number of samples (107 genes and 306 samples in this study), making data normalization a critical step for robust analysis. Using EDDA, we explored the impact of a range of normalization approaches, including the one recommended in the NanoString data analysis guide (NanoString, WA, USA). As shown in Figure 6b, the coefficient of variation (COV) of a panel of 6 housekeeping genes *(ACTG1, ACTB, EIF4G2, GAPDH, RPLP0* and *UBE2D4*) is significantly lower, as expected, when the data is properly normalized and, in particular, this is the case when using mode-normalization which produces the lowest average COV across all the methods tested. We then investigated the effect of normalization on the power to discriminate between ER positive and negative breast cancers using a panel of 8 known ER signature genes^30^. Not surprisingly, the ability to distinguish ER positive and negative breast cancers improves significantly with proper normalization with mode-normalization providing the largest F-score (see **Methods**) among all the approaches tested (Figure 6c).

In our third case study, we critically assessed the analysis done in a recent Metagenomic study that looked into the association of markers in gut microflora with type 2 diabetes in Chinese patients^31^. Due to the complexity of the microbial community in the gut, the authors reported that they were able to assemble more than 4 million microbial genes overall. Correspondingly, since on average ∼20 million paired-end reads were generated per sample, the sequencing done here is expected to provide shallow coverage of the gene set, on average. The study involved a large number of cases (71) and controls (74) and the Wilcoxon test was used to identify differentially abundant genes in a case-control comparison. Another batch of 100 cases and controls was then used to validate biomarkers identified from the first batch. We used EDDA to generate virtual microbiome profiles and assessed the performance of the Wilcoxon test in this setting, in addition to the default panel of DATs (Table 2). EDDA analysis revealed that the Wilcoxon test was likely to have been too conservative in this setting and could have been improved upon using DATs like Metastats, which was designed for Metagenomic data and edgeR, which is commonly used for RNA-seq analysis (Figure 6d). In addition, while increased coverage is likely to improve the ability to detect true differences in the microbiome, the gains are expected to be relatively modest (for edgeR, ∼1% increase in AUC with 10-fold increase in sequencing; Figure 6d). Correspondingly, despite the shallow coverage employed in this study, it is likely to have captured a significant fraction of the biomarkers that could have been determined with more sequencing. In contrast, increasing the number of replicates is likely to have markedly improved the ability to detect true differences in the microbiome, (with edgeR, ∼7% increase in AUC by doubling the number of replicates Figure 6d). Keeping sequencing cost fixed and using 300 replicates and 5 reads per gene is thus expected to boost AUC from 0.73 (using the Wilcoxon test) in the study to 0.97 (using edgeR; Figure 6d) and sensitivity from 32% to 86% at 5% FDR.

Based on this, we reanalyzed cases and controls from the first batch in this study to identify an additional 37,664 differentially abundant genes (17% increase) using edgeR, of which a greater fraction (27% increase over the original study) were also validated in the second batch of samples (**Supplementary Table 2**). The newly identified genes highlighted previously missing aspects of the role of the microbiome in Type 2 diabetes including the identification of 24 additional gene families enriched in differentially abundant genes (**Supplementary Table 3**). In particular, this analysis detected two bacterial genes identified as multiple sugar transport system substrate-binding proteins as being abundant in cases vs controls (**Supplementary Figure 11**), as well as the enrichment of two new families for multiple sugar transport system permease proteins (K02025 and K02026). Strikingly, the newly detected genes also enabled construction of an improved microbiome-based predictive model for Type 2 diabetes (AUC of 0.96 vs 0.81 in the original study; see **Methods** and **Supplementary Figure 11**), based on the selection of 50 marker genes (see **Methods** and **Supplementary Table 4**), highlighting that improved differential abundance analysis based on informed choices using EDDA can significantly impact major conclusions from a study.

## Discussion

The case studies highlighted in the previous section are not unique in any way and point to a general trend in current design of high-throughput experiments, where commonly used rules of thumb lead to suboptimal designs and poor utilization of precious experimental resources. Considering that the market for sequencing based experiments alone is currently in the billions of dollars, savings in research budgets worldwide would be substantial with even a modest 10% improvement in study design. On the other end, with a fixed budget, optimizing study design can ensure that key insights are not missed. In particular, in many scenarios where either (a) effect sizes are small and fold-changes are marginal or (b) large effects on a few entities mask subtle effects on other entities or (c) the goal is to understand coordinated processes such as cellular pathways through enrichment analysis^32^, loss in sensitivity or precision due to unguided experimental choices can be detrimental to the study. The use of a personalized-benchmarking tool such as EDDA provides a measure of insurance against this.

With the recent, dramatic expansion in the number of high-throughput applications (largely based on DNA sequencing) as well as end-users (often non-statisticians), differential abundance testing is now frequently done by non-experts in settings different from the original benchmarks for a method. This can make it difficult to determine if a particular analysis was appropriate or lead to incorrect results. One possible approach that could account for this is to use multiple DATs to get a consensus set (also available as an option on the EDDA web-server) but this can result in overly conservative predictions. For example, in a recent analysis of RNA-seq data from two temporally-separated mouse brain samples using edgeR and DESeq (with default parameters), we found that the intersection of differentially expressed genes (at 10% FDR) contained less than 10% of the union. Breakdown of the results showed that while edgeR was primarily reporting up-regulated genes (998 out of 1189), DESeq was largely reporting down-regulated genes (875 out of 878), with no indication as to which analysis was more appropriate. EDDA simulations and analysis were then used to clarify that results from edgeR were more reliable here (FPR of 3.8% vs 9.2% at 10% FDR) and could be improved further using mode-normalization (FPR of 1.5%). Furthermore, the bias towards detecting up or down-regulated genes was intrinsic to the tests here (not affected by normalization as we originally suspected) and hence reporting the union of results was more appropriate. Examples such as this are not uncommon in the analysis of high-throughput datasets and experimental design tools such as EDDA can help provide informed answers to researchers.

We hope the results in this study serve to further highlight the still under-appreciated importance of proper normalization for differential abundance analysis with high-throughput datasets^20, 33, 34^. Normalization based on mode-statistics provides an intuitive alternative to existing approaches, exhibiting greater robustness to experimental conditions in general, >20% improved AUC performance in some conditions, as well as the ability to detect cases where proper normalization may not be feasible.

EDDA was designed to provide an easy-to-use and general-purpose platform for experimental design in the context of differential abundance analysis. To our knowledge, it is the first method that allows users to plan single-cell RNA-seq, Nanostring assays and Metagenomic sequencing experiments, where the larger number of samples involved could lead to important experimental tradeoffs. The combination of model-based and model-free simulations in EDDA allows for greater flexibility and, in particular, we provide evidence that the commonly used Negative Binomial model may not be appropriate for single-cell RNA-seq, but a model-free approach (leveraging on a 96 cell dataset generated in this study) is better suited. Model-free simulations using EDDA can thus serve as a basis for refining new statistical tests and clustering techniques for single-cell RNA-seq. Note that a common assumption in EDDA and most statistical testing packages is that deviations from the multinomial model due to experimental biases can be corrected for and hence these issues were ignored in this study^14, 35^.

The basis for EDDA is a simple *simulate-and-test* paradigm as the diversity of statistical tests precludes more sophisticated approaches (e.g. deriving closed-form or numerical bounds on expected performance). Given the simplicity of this approach, it is even more surprising that the field has until now relied on rules of thumb. In light of this, the main contribution of this work should be seen as the demonstration that significant variability can be observed across all experimental dimensions and, therefore, lack of experimental design tailored to a particular application setting can lead to substantial wastage of resources and/or loss of detection power. We hope that the availability of EDDA through an intuitive, easy-to-use, point-and-click web-based interface will thus encourage a wide-spectrum of researchers to employ experimental design in their studies.

## Methods

### Single-cell Library Preparation and Sequencing

ATCC^®^ CCL-243™ cells (*i.e.* K562 cells) were thawed and maintained following vendor’s instructions using IMDM medium (ATCC^®^ 30-2005™) supplemented with 10% FBS (GIBCO^®^ 16000-077™). The cells were fed every 2 to 3 days by dilution and maintained between 2 × 10^5^ and 1 × 10^6^ cells/ml in 10 to 15 ml cultures kept in T25 flasks placed horizontally in an incubator set at 37 °C and 5% CO2. Cells were slowly frozen two days after feeding at a concentration of 4 million cells per ml in 100 µl aliquots of complete medium supplemented with 5% DMSO (ATCC^®^ 4X). The cryo-vials containing the frozen aliquots were kept in the vapor phase of liquid nitrogen until ready to use. On the day of the C1™ experiment, a 900 µl aliquot of frozen complete medium was thawed and brought to room temperature. The cryo-vial was retrieved from the cryo-storage unit and placed in direct contact with dry ice until the last minute. As soon as the cryo-vial was taken out of dry ice, the cells were thawed as quickly as possible at a temperature close to 37°C (in about 30 seconds).

The room temperature complete medium was slowly added to the thawed cells directly in the cryo-tube and mixed by pipetting four to five times with a 1000 µl pipette tip. This cell suspension was mixed with C_1_™ cell suspension reagent at the recommended ratio of 3:2 immediately before loading 5 µl of this final mix on the C_1_™ IFC. The C1 Single-Cell Auto Prep System (Fluidigm) was used to capture individual cells and to perform cell lysis and cDNA amplification following the chip manufacturer’s protocol for single-cell mRNA-seq (PN 100-5950). Briefly, chemistry provided by the SMARTer Ultra Low RNA Kit (Clontech) was used for reverse transcription and subsequent amplification of polyadenylated transcripts using the C1™ script 1772x/1773x. After harvest of the amplified cDNA from the chip, 96- way bar-coded Illumina sequencing libraries were prepared by tagmentation with the Nextera XT kit (Illumina) following the manufacturer’s protocol with modifications stated in Fluidigm’s PN 100-5950 protocol. The 96 pooled libraries were 51-base single-end sequenced over 3 lanes of a Hi-Seq 2000.

Raw reads for all libraries are available for download from NCBI using the following link: http://www.ncbi.nlm.nih.gov/bioproject/238846.

### Simulation of Count Data in EDDA

*Abundance Profile* (**AP**): When provided with sample data, EDDA uses the entity count and the sample abundance profile from the data (when multiple samples are provided, the counts are aggregated to get an average frequency profile) to do simulations. Users can also explicitly provide a profile or choose from among pre-defined profiles including **BP** (for BaySeq Profile; the profile used in simulations by Hardcastle et al.^15^), **HBR** (a profile derived from a “human brain reference” dataset^36^) and the profile from **Wu et al**^23^. In order to simulate with entity counts that differ from the original profile, EDDA allows users to sample entities (without replacement for subsampling and with replacement for over-sampling) from the middle 80% (entities are ordered by relative abundance), top 10% and bottom 10% independently. This procedure is designed to maintain the dynamic range of the original profile. In addition, to avoid working with entities with very low counts, EDDA allows users to filter out those with counts below a minimum threshold for all replicates (default of 10).

*Perturbation Profile:* If EDDA is provided with sample data under two conditions then the profile of differential abundance seen there is assigned to genes by keeping the relationship of mean expression and fold change. Specifically, EDDA applies a DAT (DESeq and FDR-cutoff of 5% by default, after mode normalization) to the sample data to identify differentially abundant entities and their corresponding fold-changes (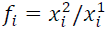 where 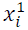 and 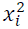 are the mean relative abundance of entity i under the first and second condition respectively). Given *d* differentially abundant entities, with fold-changes *f*1 to *f*d, a set of *d* entities from the first condition are perturbed by these fold-changes to obtain the abundance profile for the second condition (i.e. 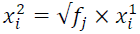 and 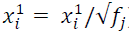) while retaining the correspondence between mean expression level and fold change. In addition, to account for undetected entities an additional fraction of entities is randomly selected (from those that fail the FDR cutoff and with fold-change > 1.5) and their observed fold-changes used to perturb abundance profiles as before. The fraction of entities was determined to ensure that the overall count matched the expected number of differentially abundant entities in the dataset (estimated from the expected number of true positives at each gene’s FDR level). In the absence of sample data, EDDA also allows users to specifically set the percentage of entities with increased and decreased entity counts (**PDA**), ranges for the fold-changes (**FC**) and the distribution to sample fold-changes from (e.g. Log-normal, Normal or Uniform).

*Simulation Model* (**SM**): The default model for simulations in EDDA is the **Full** model where the mean abundance for each entity (under each condition) and the dispersion value provided (**SV**) is used to compute means for the replicates using a Negative Binomial distribution (this is done emulating the procedure in baySeq^15^ where each entity has a dispersion sampled from a gamma distribution). When sample data is available, EDDA estimates dispersion values by using the procedure in DESeq (alternately, edgeR) to fit the empirical distribution. Entity abundances for each replicate are then normalized to get a frequency vector (that sums to 1) to simulate count data from a Multinomial distribution (where the total count is sampled from Uniform[0.9 x ND, 1.1 x ND] and ND is the number of data-points specified by the user). EDDA also allows users a simpler **Negative Binomial** model (**NB**) where the counts are directly obtained from the Negative Binomial sampling described above and a **Multinomial** model where the abundance profile (normalized to 1) is used to directly simulate from the Multinomial distribution.

*Model-Free Approach:* To create **Model-Free** simulations, EDDA requires sample data as input (ideally with many replicates and data-points). For RNA-seq, single-cell RNA-seq and Metagenomic simulations, EDDA is packaged with sample datasets discussed in this study to be used as input. If enough replicates are available in the sample dataset (i.e. greater than NR × desired number of simulations), EDDA subsamples counts from the entity in the sample dataset with the closest average count and scales counts as needed. To simulate more replicates than the number available in input data, EDDA groups entities according to their average count to sub-sample entity counts. This approach was validated using RNA-seq data^23^ where more than 90% of genes had similar expression variability as the 10 closest genes (in terms of average count; Kolmogorov–Smirnov test *p*-value < 0.05) as opposed to 2% of genes in the case of random groupings. After simulating variability in counts across replicates using the model-free approach, EDDA also provides users the option to convert counts back to a relative abundance profile for multinomial sampling of counts with a desired number of data-points.

### Mode Normalization

In principle, the ideal normalization factor for detecting differential abundance would be based on counts for an entity that is not differentially abundant (or the sum of counts for all such entities, see Figure 5). The idea behind **mode-normalization** is to identify such entities under the assumption that non-differentially-abundant entities will tend to have similar un-normalized fold changes (**UFCs**, computed as ratio of average counts across conditions). In methods such as DESeq, a related idea is implemented using a quantity called the size-factor (= ratio of observed count to a pseudo-reference sample computed by taking the geometric mean across samples) and by taking the median (or upper-quartile) size factor under the assumption that it would typically come from a non-differentially-abundant entity.

Mode-normalization in EDDA is based on calculating UFCs for all entities and determining the approximate modes for their empirical distribution (**Supplementary Figure 10**). Specifically, we used a kernel density estimation approach^37^ to smooth the empirical distribution and to compute local maxima for it. In cases where the number of maxima is not as expected (i.e. 3, corresponding to entities with decreased, unchanged and increased relative abundance), the bandwidth for smoothing was decreased as needed (starting from 0.5, in steps of 0.02, till the number of maxima is as close to 3 as possible). If the final smoothed distribution was uni- or tri-modal then the mode in the middle (presumably composed of non-differentially-abundant entities) was chosen and the normalization factor was calculated from the geometric mean of 10 entity counts around the mode. For bimodal distributions, selecting the correct mode is potentially error-prone and we flag this to the user, picking the mode with the narrowest peak (as given by the width of the peak at half the maximum value) and calculating the normalization factor as before.

### Parameter/DAT settings and EDDA extensions

EDDA is designed to be a general-purpose experimental design tool (that is easily extendable due to its implementation in R) and correspondingly it provides significant flexibility in user settings. In addition, we investigated the question of which parameter values are typically seen in common applications (e.g. RNA-seq, Nanostring analysis and Metagenomics) and used these to guide the evaluations presented in this study as detailed below.

For RNA-seq experiments, ECs in the range 1000 (microbial genomes) to 30000 (mammalian genomes) are common with NR and ND in the range [1,10] and [10,10000] per entity, respectively. For Nanostring and Metagenomic experiments (species profile), EC can be significantly lower (in the range [10, 1000]) as well as significantly higher (>1 million for Metagenomic gene profile). For the RNA-seq datasets analyzed in this study, SV was found to be in the range [0.1, 0.9] (**Supplementary Table 1**) though much higher variability was seen in Metagenomic data (> 5). Abundance profiles are best learnt from pilot data but in their absence, the sample profiles provided with EDDA should serve as a useful range of proxies. For RNA-seq experiments, PDA values in the range [5%, 30%] can be expected while Nanostring and Metagenomic experiments can have even higher percentages. Fold change distributions were typically observed to be well-approximated by a Log-normal model (**Log-normal**[µ, σ]) but other models are also feasible in EDDA (e.g. **Normal**[µ, σ] and **Uniform**[a, b]).

For DATs such as DESeq, edgeR, baySeq, NOISeq and Metastats that are implemented in R, EDDA is set to call corresponding R functions, running them with default parameters and normalization options unless otherwise specified. The results in this study were obtained with the following versions of the various packages: DESeq (v1.7.6), edgeR (v2.4.6), baySeq (v1.8.3), NOISeq (version as of 20^th^ April 2011), Metastats (version as of 14 April 2009) and R (v2.14.0). For Cuffdiff, the relevant C++ code was extracted from Cufflinks (v2.1.1) and incorporated into EDDA as a pre-compiled dynamically-linked library using Rcpp^33^ (http://cran.r-project.org/web/packages/Rcpp/index.html).

In its current form, EDDA installs the DATs listed in Table 2 by default. In addition, EDDA is designed to support the easy integration of new DATs and a step-by-step guide to do so (with the Wilcoxon test used here as an example) is provided as part of the package (see **Supplementary Text**). EDDA is also designed to be extendable in terms of simulation models and a guide for this is also provided in the installation package (see **Supplementary Text**).

### EDDA modules

For expert users the full functionality of EDDA and mode-normalization are available in a package written in the statistical computing language R that can be freely downloaded from public websites such as SourceForge and Bioconductor. In addition, to enable easy access for those who are unfamiliar with the R environment, we designed web-based modules that encapsulate typical use cases for EDDA (http://edda.gis.a-star.edu.sg; also see Figure 1a) including modules for:

a. **Differential Abundance Testing**: This module is meant to enable users to easily run a panel of DATs on any given dataset, to assess the variability of results across DATs, compute the intersection and union of these results and correspondingly select a more robust or comprehensive set of calls for downstream analysis. The assumption here is that a user has already generated all their data and would like a limited comparison of results from various DATs.
b. **Performance Evaluation**: The purpose of this module is to allow users to evaluate the relative performance of various DATs based on the characteristics of their experimental setting. A salient feature of this module is that users can adjust the stringency thresholds for the DATs and immediately assess the impact on performance, without re-running the DATs. The expected use case for this module is when users have pilot data and would like to do a systematic evaluation of the DATs.
c. **Experimental Design**: This module allows users to specify desired performance targets and the range of experimental choices that are feasible, to identify combinations that can meet the targets as well as the appropriate DATs that can be used to achieve them. Ideally, users in the planning stages of an experiment would use this module to optimize their experimental design.

### Pre-processing of RNA-seq datasets

Count data for RNA-seq datasets in ENCODE (for GM12892 NR = 3, MCF7 NR = 3 and h1-hESC NR = 4) were obtained directly from http://bitbucket.org/soccin/seqc. Count data for Pickrell et al^38^ (NR = 69) was obtained from (http://bowtie-bio.sourceforge.net/recount/). Reads from each library of the K562 single cell RNA-seq dataset were mapped uniquely and independently using TopHat^39^ (version 2.0.7) against the human reference genome (hg19). Raw counts for each gene were then extracted using Human Gencode 19 annotations and htseq-count (http://www-huber.embl.de/users/anders/HTSeq/doc/overview.html).

### Single-cell RNA-seq and Metagenomic Analysis

RNA-seq count data from single-cell experiments in Ramskold et al^29^ was obtained from Supplementary Table 4 in the manuscript (15 thousand genes and 10 samples - 6 putative circulating tumor cells and 4 from the melanoma cell line SKMEL5; RPKM values were converted back to raw counts). Metagenomic count data from the study by Qin et al^31^ was obtained from www.gigadb.org (1.14 million genes and 145 samples). The fit of the metagenomic count data to the Negative Binomial distribution was assessed using a Kolmogorov–Smirnov test, where < 0.1% of the genes failed the test at a *p*-value threshold of 0.01. Characteristics of both datasets were learned by EDDA and used to generate simulated datasets (Model-Free for single-cell and with the Full Model for metagenomic data).

To build a predictive model for type 2 diabetes from the data in Qin et al^31^, we followed the procedure described there to identify 50 marker genes from the top 1000 differentially abundant genes (based on edgeR *p*-values) by employing the maximum relevance minimum redundancy (mRMR) feature selection framework^40^. The identified marker genes were combined into a “T2D index” (= mean abundance of positive markers - mean abundance of negative markers) for each sample, which was then used to rank samples and compute ROC curves as in the original study.

### Nanostring Analysis

Nanostring count data was obtained from an in-house preliminary study of prognostic and predictive gene signatures for breast cancer (manuscript in preparation). Briefly, expression levels of 107 genes of interest from 369 patients in different stages of breast cancer and with known estrogen receptor (ER) alpha status and clinical outcomes were quantified using the NanoString nCounter System (NanoString, WA, USA). The raw data was normalized by different generally applicable methods (e.g. median-normalization as implemented in DESeq, mode-normalization from the EDDA package and UQN as implemented in edgeR; see Table 2) as well as the recommended standard from the NanoString data analysis guide (normalized by positive and negative controls, followed by global normalization). Note that as this dataset has multiple categories we extended the standard two-condition version of mode-normalization by randomly labelling samples as controls or cases to identify the top 10 genes that are consistently chosen. In order to measure the impact of normalization on the ability to separate patients based on their ER status a standard F-score was calculated, as the ratio of between-group variance to within-group variance of mean counts for the 8 ER signature genes (formally F-score = *F*_*Between*_/*F*_*Within*_, where *F*_*Between*_ = |*X*_1_|(*E*(*X*_1_) − *E*(*X*))^2^ +|*X*_2_|(*E*(*X*_2_) − *E*(*X*))^2^ and *F*_*within*_ = (|*X*_1_|*Var*(*X*_1_) + |*X*_2_|*Var*(*X*_2_))/(|*X*_1_|+|*X*_2_| − 2) and *X* = *X*_1_ ∪ *X*_2_, for mean counts *X*_1_ and *X*_2_ in the two groups).

## Availability

As an open-source R package at https://sourceforge.net/projects/eddanorm/ or http://www.bioconductor.org/packages/devel/bioc/html/EDDA.html and as web-modules at http://edda.gis.a-star.edu.sg.

## Acknowledgements

We would like to thank Shyam Prabhakar, Swaine Chen, Denis Bertrand, Sun Miao, Li Yi and Chng Kern Rei for providing valuable feedback on drafts of the manuscript, Christopher Wong for sharing the Nanostring dataset with us and Lili Sun, Naveen Ramalingam and Jay West for technical assistance in generating the single-cell RNA-seq dataset.

### Funding

This work was done as part of the IMaGIN platform (project No. 102 101 0025), supported by a grant from the Science and Engineering Research Council as well as IAF grants IAF311009 and IAF111091 from the Agency for Science, Technology and Research (A*STAR), Singapore.

### Authors Contributions

NN and CKHB initiated the project. LH and CKHB implemented EDDA with additional inputs from LJ. LH designed and implemented mode-normalization with inputs from NN. PR coordinated the single-cell RNA-seq data generation. LJ conducted the single-cell and metagenomic analysis. LH conducted the nanostring analysis. NN, LH, LJ and CHKB wrote the manuscript with inputs from all authors.

### Competing Financial Interests

None

